# Can structure predict function at individual level in the human connectome?

**DOI:** 10.1101/2023.12.11.568977

**Authors:** Lars Smolders, Wouter De Baene, Geert-Jan Rutten, Remco van der Hofstad, Luc Florack

## Abstract

Several studies predicting Functional Connectivity (FC) from Structural Connectivity (SC) at an individual level have been published in recent years, each promising increased performance and utility. We investigated three of these studies, analyzing whether the results truly represent a meaningful individual-level mapping from SC to FC. Using data from the Human Connectome Project shared accross the three studies, we constructed a predictor by averaging FC of training data and analyzed its performance in the same way. In each case, we found that this group average FC is an equivalent or better predictor of individual FC than the predictive models in terms of raw prediction performance. Furthermore, we showed that additional analyses performed by the authors of the three studies, in which they attempt to show that their predicted FC has value beyond raw prediction performance, could also be reproduced using the group average FC predictor. This makes it unclear whether any of the three methods represent a meaningful individual-level predictive model. We conclude that either the applied methods are not appropriate for the data, that the sample size is too small, or that the data itself does not contain sufficient information to learn a mapping from SC to FC e.g. due to the amount of noise in MRI measurements. We advise future individual-level studies to always explicitly report their results in comparison to the performance of the group average, and carefully demonstrate that their predictions contain meaningful individual-level information. Finally, we believe that investigating alternatives for the construction of SC and FC may improve the chances of developing a meaningful individual-level mapping from SC to FC.

## Introduction

Understanding how anatomical connections in the brain give rise to functional interactions, which in turn support the cognitive functioning of individuals, is one of the major goals in neuroscience. In recent years, these ideas have been conceptualized in terms of networks consisting of connections between regions of interest (Bullmore & Sporns, 2009; Honey et al., 2009; Sporns et al., 2005). Structural connections are represented by structural connectivity (SC), which quantifies the strength of white matter tracts connecting each pair of brain regions (Hagmann et al., 2008; Sporns et al., 2005; van den Heuvel & Hulshoff Pol, 2010). Functional interactions are represented by functional connectivity (FC), which expresses the degree of co-activation between pairs of regions (Beckmann et al., 2005; Damoiseaux et al., 2006). There exists a substantial body of work that attempts to predict FC from SC, with approaches such as biophysical models and network communication models. However, the performance achieved by these models is modest, and usually only reported at a group level (e.g., Avena-Koenigsberger et al., 2018; Goñi et al., 2014; Mišić et al., 2016; Seguin et al., 2018; Wang et al., 2019). For clinical applications, such as neurosurgical planning, we must take into account that the organization of each patient’s brain network can vary strongly, especially when considering brain tumor patients (Almairac et al., 2021; Harris et al., 2014; Jütten et al., 2020; Maesawa et al., 2015; Park et al., 2016). It is therefore essential that predictions of function from structure are accurate at an individual level if we wish to implement these methods in clinical practice.

In the past years, several studies have attempted to predict function from structure at an individual level (Becker et al., 2018; Benkarim et al., 2022; Cummings et al., 2022; Deslauriers-Gauthier et al., 2020; Neudorf et al., 2022; Sarwar et al., 2021), generally based on machine learning techniques. We closely examined three of these studies, which demonstrated the most promising results on large and diverse data sets, to determine whether their methods are suitable for further development in a clinical setting. Each of the three studies followed the existing literature, calculating FC as the Pearson correlation between the activity levels over time in each brain region, while calculating SC from the number of white matter fibers as found by tractography connecting each pair of brain regions. The studies then attempted to predict FC from SC in a number (*n* between 250 and 1000) of healthy subjects from the Human Connectome Project (HCP), which allows us to compare their results. Two of the studies also considered supplementary data sets.

In the following, we briefly summarize the three studies we examined. After this, we show that we can reproduce their results using the group average FC with added random noise, which does not contain individual information. From this we conclude that the authors of the three studies have not demonstrated that the learned individual-level mappings preserve individual information. This might indicate that either individual-level methods as they are currently applied in the task of predicting function from structure are not yet suitable, or that the data they are applied to does not contain sufficient information to learn a mapping from.

### Predicting individual FC from SC: a summary

In **Structure-function coupling in the human connectome: A machine learning approach,** Sarwar and colleagues (2021) trained a 10-layer fully connected neural network with a loss function designed to preserve inter-individual variation in FC, ensuring that predictions did not converge to a group average. This loss function was parametrized by two parameters, γ and λ, which induced a trade-off between prediction accuracy and inter-individual variation in predictions. The final tuned model achieved a correlation between predicted FC (pFC) and empirical FC (eFC) of 0.9 ± 0.1 on a group level and 0.55 ± 0.1 on an individual level, improving upon the performance of earlier work using biophysical and network communication models. On the data used in the supplementary materials, the performance was even higher, reaching a correlation of 0.7 ± 0.1 at an individual level. The differing performance is probably due to the use of different pre-processing and connectivity mapping methods. The authors simultaneously demonstrated that the model did not predict a group average by explicitly showing that the inter-subject variance in pFC is similar to that in eFC. Furthermore, the authors showcased the usefulness of individual-level pFC, in which they predicted variation in cognitive performance from pFC. The results suggest that predicting cognition from pFC works almost as well as predicting cognition from eFC, achieving a correlation between predicted and actual cognitive performance of 0.33 using eFC and 0.29 using pFC. This further supports the conclusion that the machine learning model preserved relevant individual information contained in SC, and did not simply output some averaged FC that correlates well with the whole data set.

In **Structure can predict function in the human brain: a graph neural network deep learning model of functional connectivity and centrality based on structural connectivity**, Neudorf et al. (2022) used a deep learning model more directly suited to the task. Leveraging the network structure present in SC and FC, they applied a Graph Neural Network (GNN) model, trained on the same data used by Sarwar et al. Furthermore, they explicitly trained another model to predict the degree centrality, eigenvector centrality and PageRank centrality of nodes in FC, which are known to be related to age and sex differences (Zuo et al., 2012) and to various pathologies (Chen et al., 2015; Deng et al., 2019). The results were presented in terms of variance explained, where the authors show that pFC explains 56% of the variance in eFC i.e. R^2^ = 0.56. Since this is a univariate regression, we can convert to correlation by taking the square root, which yields a correlation of R = √0.56 = 0.75. While the authors suggested that this means that their method outperforms the methods by Sarwar et al. in terms of raw performance, Sarwar et al. reported their correlation as an average of correlations per individual, rather than the single correlation of all individuals’ eFC and pFC concatenated. Therefore, these numbers are not comparable. Besides raw prediction performance, network centrality measures calculated from pFC also correlated with centrality measures calculated from eFC at individual level, explaining 81% of the variance in degree centrality. The authors thus demonstrated that the model learned a mapping from SC to FC that also preserved the network structure of FC.

Finally, in **A Riemannian approach to predicting brain function from the structural connectome**, Benkarim and colleagues (2022) developed a more complex framework for FC prediction, using a generalized diffusion model on the SC network to simulate communication dynamics. This model allows the authors to explicitly model the “walk length” that is used, which represents the number of connections that are used by information propagating along the structural network. The hyperparameters were fitted on a small subset of data (n = 26) and validated on 250 subjects from the HCP data set. The presented results improved upon the results of Sarwar et al. in terms of raw performance (individual correlation = 0.775 ± 0.049 on the 200-parcel Schaefer atlas), and the authors provided some additional analyses investigating in which parts of the brain FC is harder to predict. In particular, they found that their method performed better at predicting FC in the visual, somatomotor and attention regions of the brain than in the default mode, frontoparietal and limbic regions. Furthermore, predictions in the default mode and frontoparietal regions benefitted the most from using longer walks on the structural connectome. This suggests that these regions require more polysynaptic connections to function, in agreement with earlier results in the literature indicating the same dependencies (Suárez et al., 2020; Vázquez-Rodríguez et al., 2019).

As a measure of accuracy, one must be careful when considering correlation. Specifically, given some subject’s eFC, there exist an infinite number of FC profiles that have the same correlation with this eFC. This means that, by itself, an average individual-level correlation of 0.55 or 0.77 gives no guarantee that a model preserves useful information contained in an individual’s eFC. Furthermore, in the setting of predicting FC, we must consider that the group average FC is a very good predictor by itself in terms of raw performance, as identified earlier by Deslauriers-Gauthier and colleagues (2022). In the following, we show that pFC in each of the three studies does not outperform a predictor that outputs the average eFC of the training data with added random noise. Critically, this means that a predictor based on no individual information (i.e. not requiring any input) can match the performance of complex machine learning methods.

Considering this, it is important to judge the quality of results from prediction methods not only by raw performance (in terms of correlation), but also by other metrics. Each of the three considered studies analyzed some other aspect of pFC besides raw prediction performance. We show that each of these analyses can also be reproduced using the group average eFC. This means that none of the three studies sufficiently demonstrated that pFC as generated by their methods contains more individual information than a group average eFC.

## Methods

### Data set

We used data from 1000 healthy participants of the Human Connectome Project (HCP) (Van Essen et al., 2013), a data set shared across the three studies. For each subject, we obtained minimally preprocessed diffusion-weighted and resting-state functional MRI images. Preprocessing pipelines are described in (Van Essen et al., 2013).

### Construction of SC and eFC

Since the three studies used different cortical parcellations (atlases), we also used multiple atlases to control for differences in performance. Sarwar et al. presented most of their results on the Desikan-Killiany (DK) cortical parcellation (Desikan et al., 2006). Neudorf et al. presented their results for both the DK and Automated Anatomical Labeling (AAL) atlas (Tzourio-Mazoyer et al., 2002), achieving the highest performance on the DK atlas. Finally, Benkarim et al. used a functional parcellation by Schaefer et al. (Schaefer et al., 2018) and a structural parcellation based on a refinement of the DK atlas (Vos de Wael et al., 2020), attaining the best results with the Schaefer atlas. Given that the best results and most additional analyses across the three studies are based on the DK and Schaefer atlases, we decided to use the DK and 200-parcel Schaefer atlases in our analyses.

We calculated SC and eFC matrices in the same way as described by Sarwar et al. To construct SC matrices, we obtained minimally preprocessed diffusion-weighted MRI scans for each HCP subject. Details of preprocessing are described in (Van Essen et al., 2013). In short, spin-echo planar imaging was used to acquire 90 gradient directions at b-values of 1000, 2000 and 3000 s/mm^2^. The images were corrected for head motion, gradient-nonlinearity distortions and eddy currents, and subsequently normalized to standard MNI space. We performed whole-brain tractography on this data using a deterministic algorithm (as suggested in (Sarwar et al., 2019)), estimating fiber orientations with constrained spherical deconvolution (Tournier et al., 2007) using a spherical harmonic order of 8. Using a white matter mask generated from the diffusion data, dilated one voxel to account for the white and grey matter boundary, we uniformly seeded two million streamlines, propagated using the *tckgen* function in MRtrix (Tournier et al., 2019) with the *sd_stream* algorithm. Default tractography parameters were used (step size 0.1 *x* voxel size, angle threshold 9°*x* step size / voxel size, FOD threshold 0.1) and maximum streamline length was set to 400mm. This whole-brain tractogram was mapped into an SC matrix by counting the number of streamlines connecting each pair of regions in the considered atlases (DK and Schaefer). To account for order of magnitude differences in matrix entries that result from this procedure, a Gaussian resampling method was applied (Honey et al., 2009). Briefly, a value is sampled from a standard normal distribution for each matrix entry. These values are then assigned to the SC matrix in order, i.e. the smallest sampled value is assigned to the smallest SC value, the second-smallest sampled value to the second-smallest SC value, and so on until the SC matrix is fully replaced by sampled values. Finally, the SC matrix was rescaled to a mean of 0.5 and standard deviation of 0.1.

For eFC, we used two minimally preprocessed resting-state functional MRI images, one with right-to-left and one with left-to-right phase encoding, for each HCP subject. fMRI images were acquired using a multiband gradient-echo planar imaging period of 14.4 minutes, with a repetition time of 720ms and a multiband factor of 8. Subjects were instructed to keep their eyes open and fixate on a projected cross-hair on a dark background. Preprocessing is again described in (Van Essen et al., 2013). In short, the images were denoised and motion-scrubbed with ICA-FIX (Salimi-Khorshidi et al., 2014), then normalized to standard MNI space. The two images were temporally demeaned and concatenated, resulting in a single, longer, image. Activity time series were calculated for each region in the considered atlases (DK and Schaefer) by averaging the comprising voxel activity time series. eFC was then calculated as the pair-wise Pearson correlation coefficient between each region’s time series.

### Assessing prediction performance

Following Sarwar et al., we adopted a 10-fold cross-validation framework for predicting eFC. We split the data set in 10 folds of equal size, using 9 folds as training data and the remaining fold to assess performance i.e. training on 900 subjects and assessing performance on 100 subjects. Repeating this 10 times, using each fold for validation exactly once, we assessed the performance of predictors described below on the full data set. Firstly, we calculated the individual-level performance of predictors by correlating each subject’s eFC with the predictor and reporting the resulting distribution over all subjects. Secondly, we concatenated the eFC and predictors for each subject, and calculated the variance explained on the full data set. We then compared the performance of our predictors to those of Sarwar et al., Benkarim et al., and Neudorf et al. side by side.

### Noisy average eFC

Given the fact that the group average eFC is a good predictor for individual eFC (Deslauriers-Gauthier et al., 2022), we can artificially obtain the performance of any predictor with a lower or equal performance by independently adding randomly generated noise to the group average for each subject. In this procedure, we first calculated the group average eFC of the training data. Then, for each subject in the validation set, we generated random noise from a normal distribution and assigned the training group average eFC plus noise to this subject. This way, we also simulated inter-individual variability, since the added noise between any two subjects was not related. This predictor was named *noisy average FC* (navgFC). Note that navgFC does not contain subject-specific information, since the only data we used to construct navgFC was the training eFC, which did not contain the eFC or SC matrices of the subject under consideration.

### Reproducing deep learning model by Sarwar et al

In order to reproduce the analyses conducted by Sarwar et al. described below, we reconstructed the deep learning model described in their paper (i.e. ten fully connected feed-forward layers of 1024 hidden neurons, for details see (Sarwar et al., 2021)), training it to predict eFC from SC using the same loss function with γ = 0.4 and λ = 0.01 in 20000 epochs. Training was performed using a 10-fold cross validation framework, where we generated pFC by having the model predict the FC of subjects in the validation folds.

### Prediction of cognitive performance

Sarwar et al. presented additional analyses in which they attempted to predict cognitive performance of individuals from their pFC and their eFC. Using pFC as generated by the deep learning model as well as pFC generated by a biophysical model, the authors first regressed out SC in order to remove the strong linear dependence that is retained by the biophysical model. The residuals were then used to predict cognition, resulting in a correlation of 0.33 when using eFC, 0.29 when using pFC from deep learning, and 0.21 when using pFC from the biophysical model. Based on this, the authors argued that pFC retained information about individuals that could be used to predict cognition.

We conducted additional analyses regarding the predictive power of pFC. In particular, we focused on the step in which SC is regressed out of pFC, and examined whether this step unintentionally obfuscates the predictive power of pFC. We performed the analyses as described by Sarwar and colleagues, predicting the same cognition outcome from FC using lasso regression in a 10-fold cross validation. We repeated this process in 10 outer cross validation loops using different randomized folds, in order to prevent the results from depending on a specific subdivision of subjects. We differentiated between predictors with SC regressed out and predictors without SC regressed out by using the suffix “\SC”. To examine the reliance of predictive performance of pFC\SC on SC, we constructed an alternative predictor by randomly shuffling pFC between subjects (rpFC) and then regressing out the SC of the “correct” individual (i.e. when predicting cognition of subject A, we take a random other individual’s pFC and then regress out the SC of subject A). This predictor was named rpFC\SC. Furthermore, we analyzed the predictive properties of noisy average FC (navgFC), with noise tuned to capture performance and inter-individual correlations as presented in Figure2b. Finally, we used average training eFC without noise (avgFC) as predictor. Correlation coefficients and p-values were calculated by correlating the predicted cognition values with the actual cognition values of each inner cross-validation fold concatenated.

### Analyzing differences in performance across the cortex

Benkarim et al. showed how each of Yeo’s 7 brain networks (Yeo et al., 2011) contributed to the prediction errors in their pFC and how these errors depended on the walk length considered by their model. We replicated these analyses using our group average predictor by calculating the same error measure as Benkarim and colleagues (*log(1 – correlation)*) separately on each parcel of the Schaefer 200-parcel atlas. We then averaged the errors over each of Yeo’s 7 networks using the network identities provided with the atlas and finally averaged the resulting network-level errors over all subjects.

### Prediction of centrality measures

Neudorf et al. presented additional analyses in which they show that their model can also explain variance in centrality measures of eFC. The authors calculated degree centrality, eigenvector centrality and PageRank centrality of each node in both the eFC and pFC matrices. The measures were calculated on a thresholded version of the FC matrices, where a significance threshold of *p* = 0.0001 on the correlation coefficients was used (Zuo et al., 2012). The centrality measures were then z-score standardized and rescaled such that their values ranged from -1 to 1. Then, all values were concatenated for all subjects and the total variance in eFC centrality explained by pFC centrality was presented. Again adopting the 10-fold cross-validation framework, we computed the centrality measures on all eFC matrices using the DK atlas with the described procedure and averaged the cenrality values over all training subjects. These average centralies per node were then used as predictors for eFC centrality measures. The authors also trained a separate model to specifically predict centrality measures, which performed better at explaining variance in centrality measures than pFC. However, since we are only interested in the information contained in pFC, we did not consider this model in our analyses.

## Results

### Performance of group average eFC

We observed that the group average eFC matrix scores an average correlation with individual-level eFC of 0.84 using the DK atlas (see Figure 1a). Taking the group average of the training set as predictor for the validation set in each fold, we achieve a nearly identical performance distribution (Figure 1b). Similarly, we find an average correlation of 0.76 using the Schaefer atlas (Figure 1c-d).

**Figure 1:**
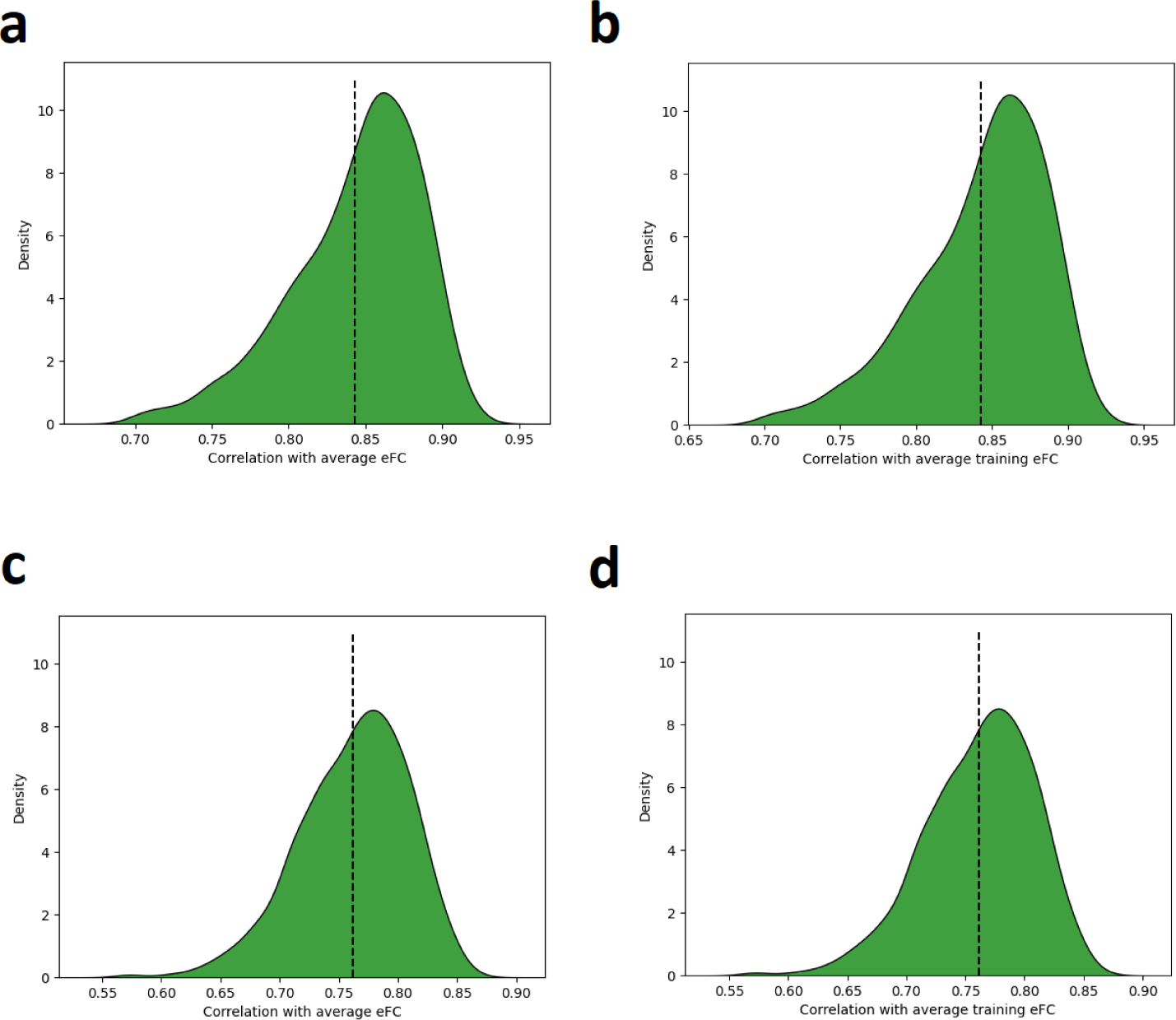
Distribution of correlations between individual eFC for each HCP subject and **a)** whole group average eFC (DK atlas); **b)** training set group-average eFC (DK atlas); **c)** whole group average eFC (Schaefer atlas); and **d)** training set group-average eFC (Schaefer atlas).

We clearly see that the performance benchmark is different for each atlas, which we should take into account when comparing performances in terms of correlations between studies.

### Performance of noisy group average eFC

Using a standard deviation of 0.1 for the noise in navgFC (chosen via manual tuning) yields correlational distributions with the same averages as the distributions presented by Sarwar et al. (Figure 2). Note that we have used plots from the supplementary material of Sarwar et al. here (Sarwar et al. (2021) Figure S1e-f), since these correlation distributions matched our own experiments on the same data, in which we found inter-individual variation in eFC (inter-eFC) to center around 0.7. The prediction performance of Sarwar et al. (Figure 2a, right-hand plot, left-most boxplot) is matched by navgFC (Figure 2b, right-most boxplot). Importantly, the average inter-individual variation in pFC (inter-pFC) of Sarwar et al. (Figure 2a, left-hand plot, red distribution) is also matched by navgFC (Figure 2b, middle boxplot).

**Figure 2:**
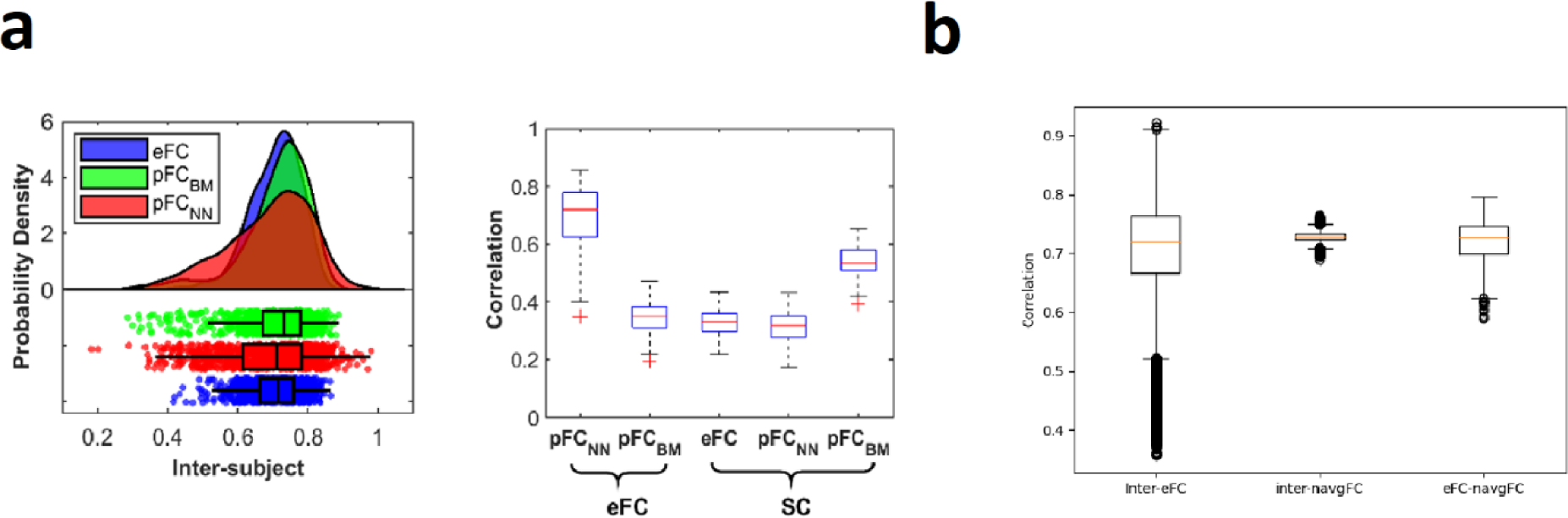
comparison of performances reported by Sarwar et al. with navgFC performance. **a)** Distributions of inter-individual variation and prediction performance as reported by Sarwar et al. in supplementary materials (adapted from (Sarwar et al., 2021) Figure S1e-f under <u>CC-BY 4.0 license</u>, no changes made). The left-hand plot shows inter-subject variance for eFC (blue) and pFC as generated by the deep learning model (red). In the right-hand plot, the first boxplot shows the prediction performance of the deep learning model. **b)** Distribution of correlations between subjects’ eFC, between 1000 generated navgFC instances, and between eFC and navgFC per subject using the DK atlas.

Since the performance reported by Benkarim et al. is nearly equal to the performance of the group average (0.775 using the 200-parcel Schaefer atlas), adding no noise (or equivalently, noise with 0 standard deviation) yields a distribution with nearly the same average (Figure 3). The results reported by Benkarim et al. are slightly better than that of the group average here (0.775 versus 0.76 respectively), but this small difference might well be attributed to minor differences in preprocessing.

**Figure 3:**
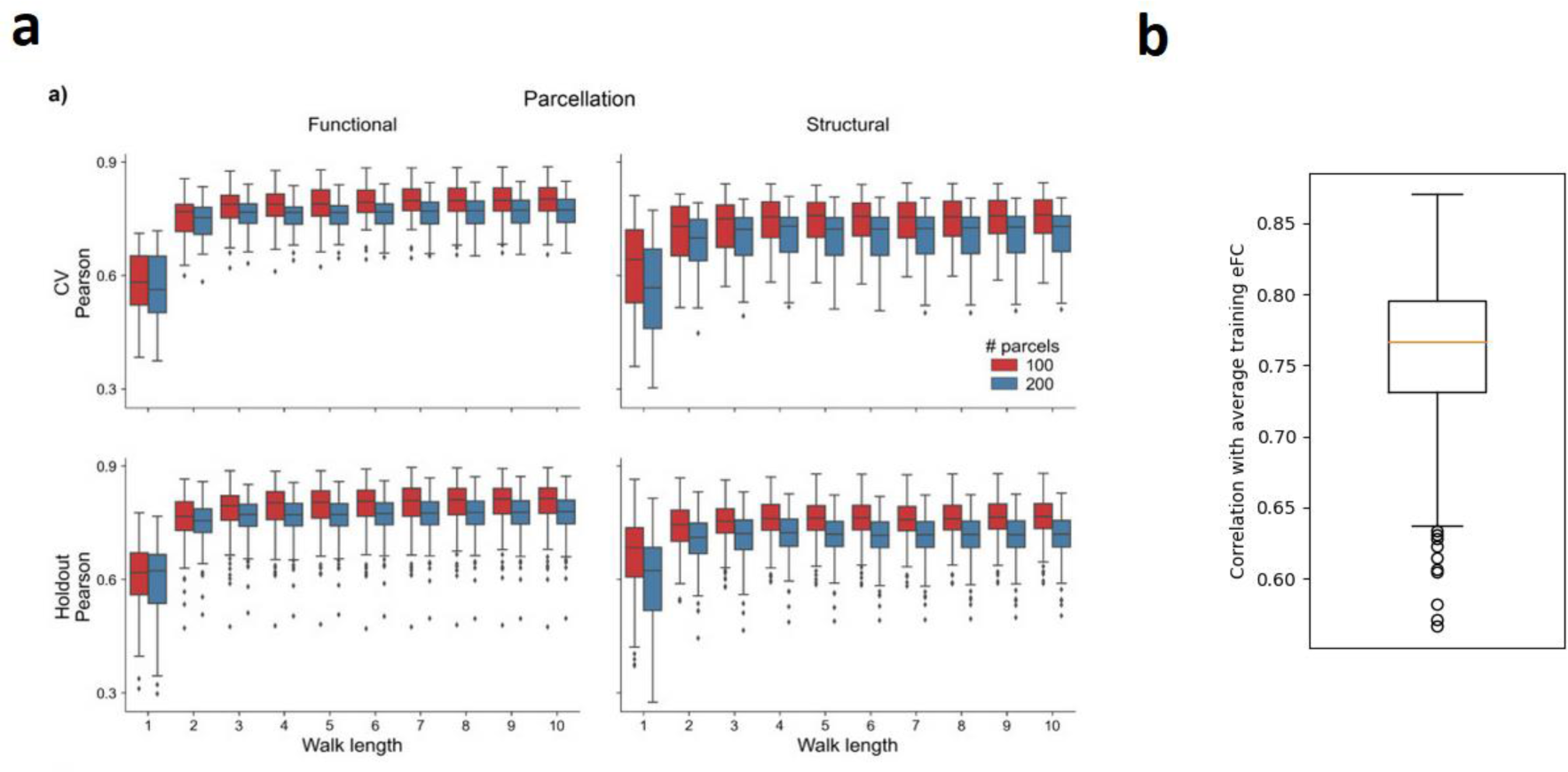
comparison of performances reported by Benkarim et al. with navgFC performance. **a)** Performance as reported by Benkarim et al. (copied from (Benkarim et al., 2022) Figure 2a under <u>CC-BY-NC-ND 4.0 license</u>) for varying walk lengths. The bottom-left plot corresponds to the 200-parcel Schaefer atlas used in our analyses. In this plot, the right-most blue boxplot corresponds to the best obtained performance. b) Performance of navgFC with zero standard deviation using the 200-parcel Schaefer atlas.

Neudorf et al. did not explicitly report distributions of performance, but rather reported performance in terms of variance explained (R^2^ = 0.56) in the whole data set using the DK atlas. If we insert the training group average (i.e. navgFC with 0 standard deviation) using the DK atlas as predictor for each subject, then the R^2^ is also almost equal to 0.56 (Figure 4).

**Figure 4:**
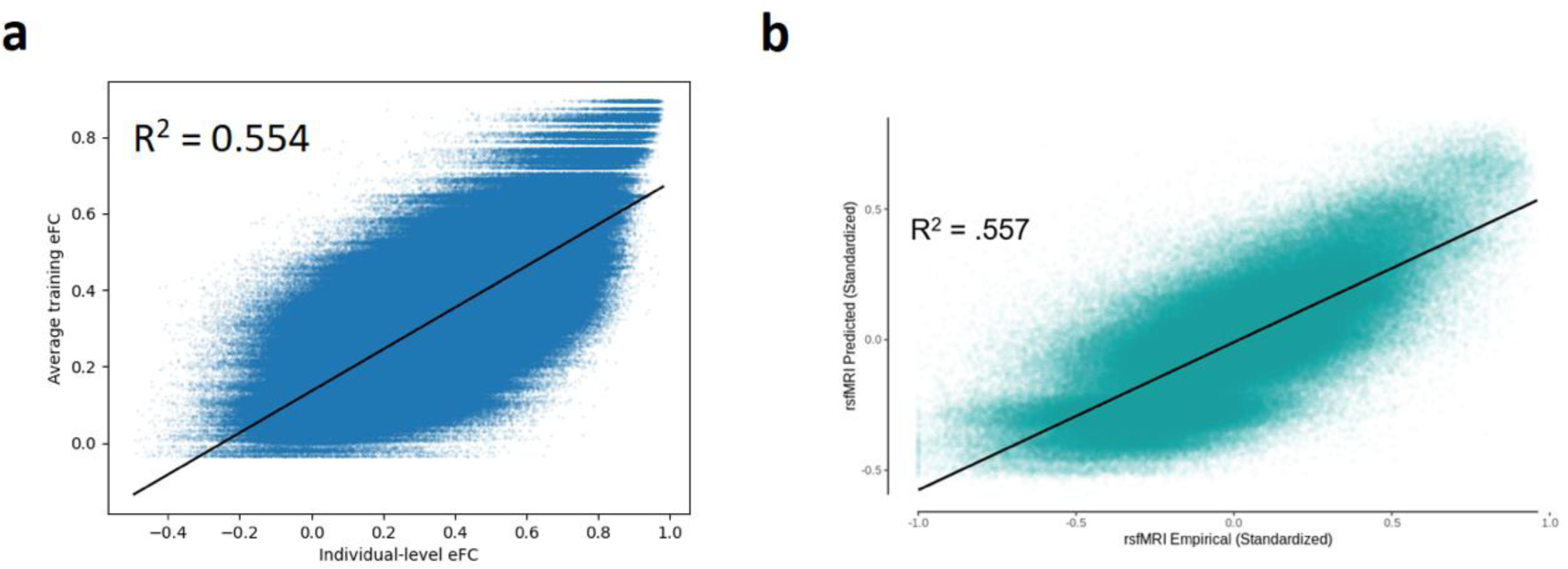
Relation between predictors and individual-level eFC for **a)** average training eFC (i.e. navgFC with zero standard deviation) using the DK atlas and **b)** pFC by Neudorf et al. (adapted from (Neudorf et al., 2022) Figure S1 under <u>CC-BY 4.0 license</u>, no changes made)

Clearly, navgFC has the same average inter-subject correlation as reported by Sarwar et al., as well as the same average performance predicting eFC in terms of correlation as all three studies when the noise is appropriately tuned.

### Preservation of individual differences

It is important to note that navgFC does not preserve inter-individual correlations (i.e. pairs of subjects with high inter-eFC may have low inter-navgFC and vice versa). In particular, since the noise is normally distributed and added independently for each subject, this correlation is exactly 0 on average. However, although Sarwar et al. showed that in their study average inter-pFC is equal to average inter-eFC, these inter-individual correlations are not preserved at an individual level in pFC either.

We generated pFC matrices for each subject using the deep learning network proposed by Sarwar et al., as described in our methods. For each pair of subjects, we calculated the correlation between their eFC matrices and their pFC matrices. Plotting each of these as data points, with inter-pFC as function of inter-eFC, yields Figure 5, where we find that the correlation between individual-level inter-eFC and inter-pFC is almost 0. Even though the distribution of inter-individual variation is conserved in the study by Sarwar and colleagues, the individual differences are not preserved at an individual level. In the studies of Benkarim et al. and Neudorf et al., the variability in predicted pFC is not quantified, which means there is no evidence that the prediction models preserve individual differences.

**Figure 5:**
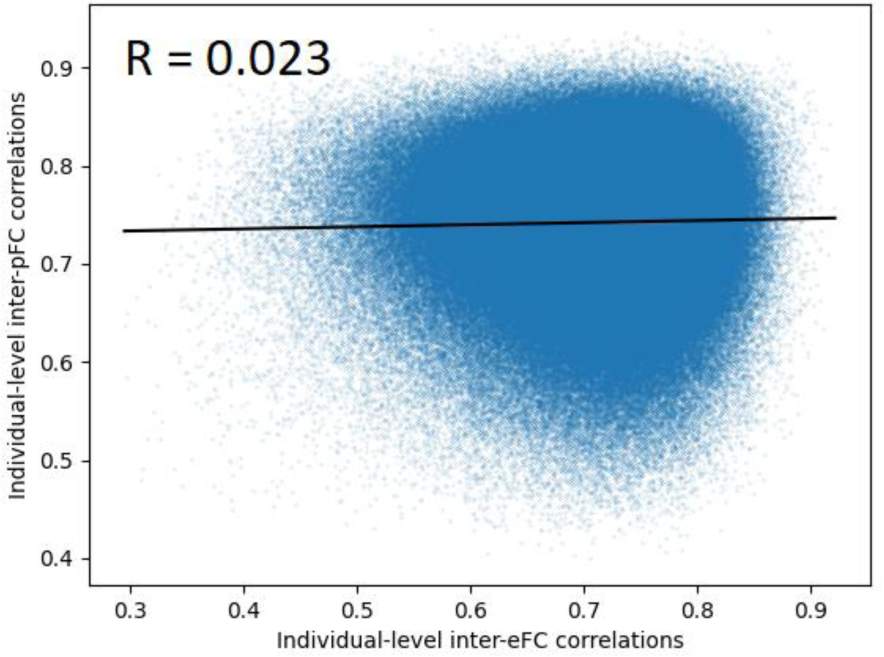
Relation between inter-eFC and inter-pFC at individual level, resulting from eFC and pFC generated by following the methods of Sarwar et al.

### Predicting cognitive performance

Again using pFC as generated by the deep learning model proposed by Sarwar et al., we found that eFC, SC, pFC\SC, rpFC\SC, navgFC\SC and avgFC\SC could significantly predict cognition, while pFC, rpFC, navgFC and avgFC could not (Figure 6). The fact that rpFC\SC can significantly predict cognition implies that it does not matter whose pFC matrix we use when predicting cognition, as long as we regress out the correct subject’s SC. Furthermore, we see that pFC by itself cannot significantly predict cognition. Finally we observe that average FC with and without added random noise can predict cognition when SC is regressed out. This means that even starting with a predictor which contains no personalized information can produce a significant predictor if SC (with personalized information) is regressed out. Therefore, we conclude that the predictive power of pFC\SC relies on the step in which SC is regressed out, rather than information retained in pFC itself.

**Figure 6:**
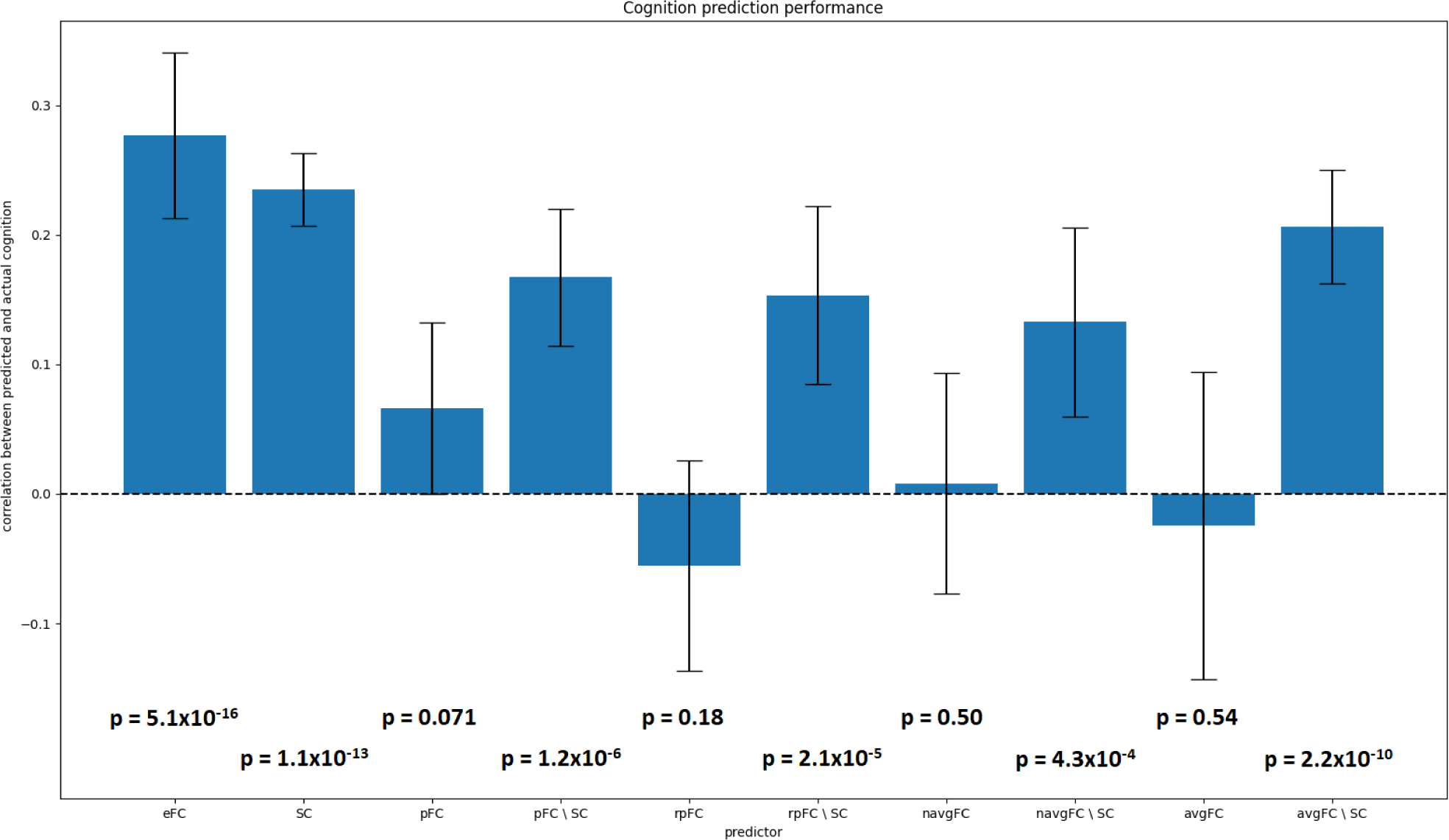
performance of several predictors in predicting cognition using Lasso regression. Error bars indicate variance of performance between outer cross validation loops.

### Differences in performance across the cortex

Benkarim et al. found that their model performed worse in predicting eFC in the default mode, frontoparietal and limbic networks than in the visual, somatomotor and attention networks of the brain. Furthermore, predictions in the default mode and frontoparietal regions benefitted the most from using longer walks on the structural connectome (Figure 7a). Inserting the group average eFC into this analysis revealed a similar pattern of errors (Figure 7b) when comparing them to the right-most data points in Figure 7a. The dependency on walk length observed by Benkarim and colleagues can likely be attributed to a general lower performance when using shorter paths. We conclude that also for this study, we cannot determine whether the model predicts something more informative than the group average.

**Figure 7:**
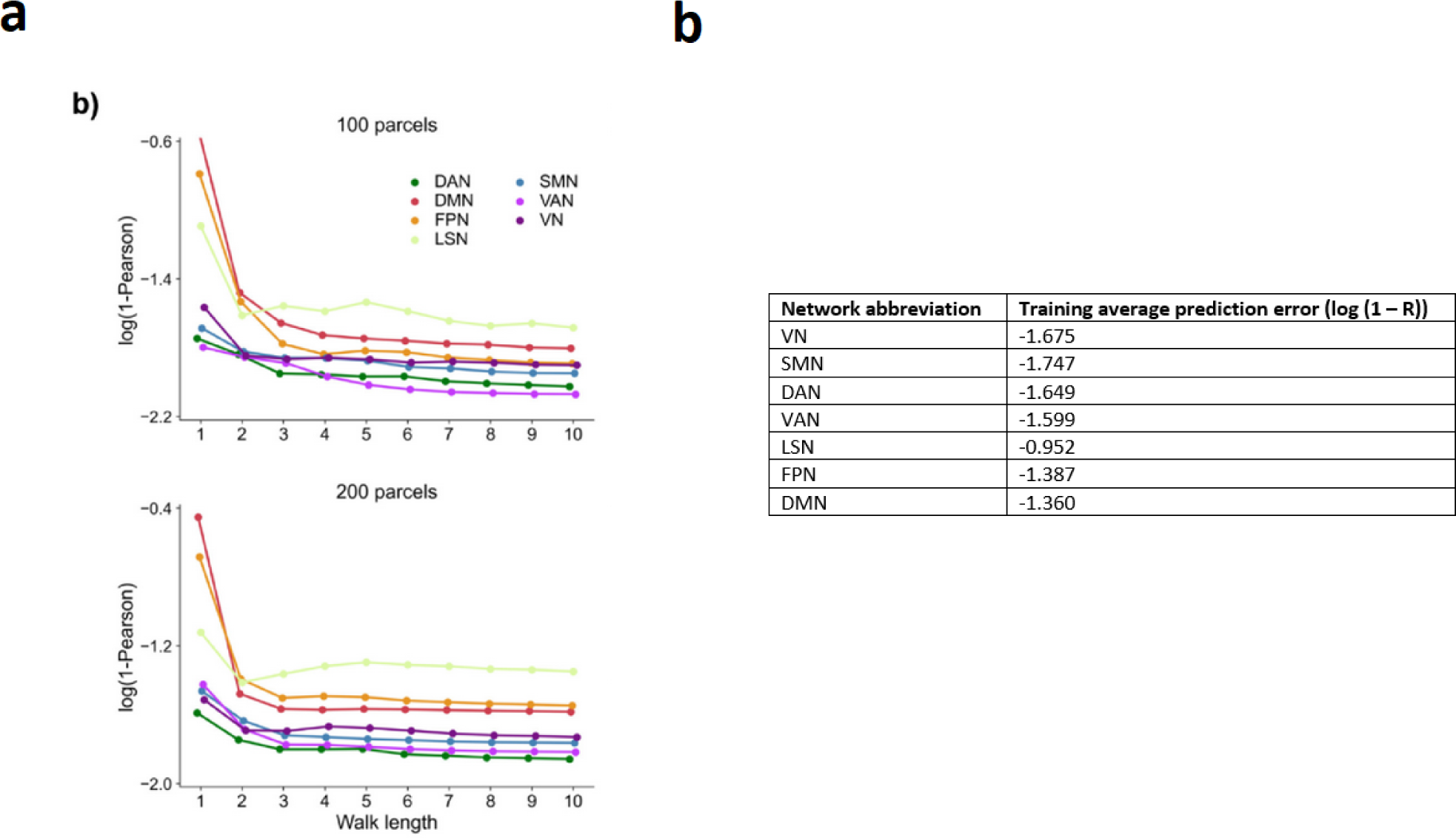
Prediction errors per Yeo’s functional network **a)** as reported by Benkarim et al. (copied from (Benkarim et al., 2022), Figure 4b under <u>CC-BY-NC-ND 4.0 license</u>) **b)** using the training average eFC as predictor

### Explaining network centrality measures in eFC

Using average centrality of each node as predictor for the centrality of each validation subject’s nodes yielded the same or better performance as reported by Neudorf et al. (Figure 8). The variance in degree centrality explained by the training group average was slightly smaller than that of pFC as reported by Neudorf et al., but this small difference can again be attributed to minor differences in preprocessing. In eigenvector and PageRank centrality, the training group average even performs much better than pfC (Figures 8b-c). From this we conclude that there is no evidence that the graph neural network presented by Neudorf et al. has learned anything other than the group average with some minor deviations.

**Figure 8:**
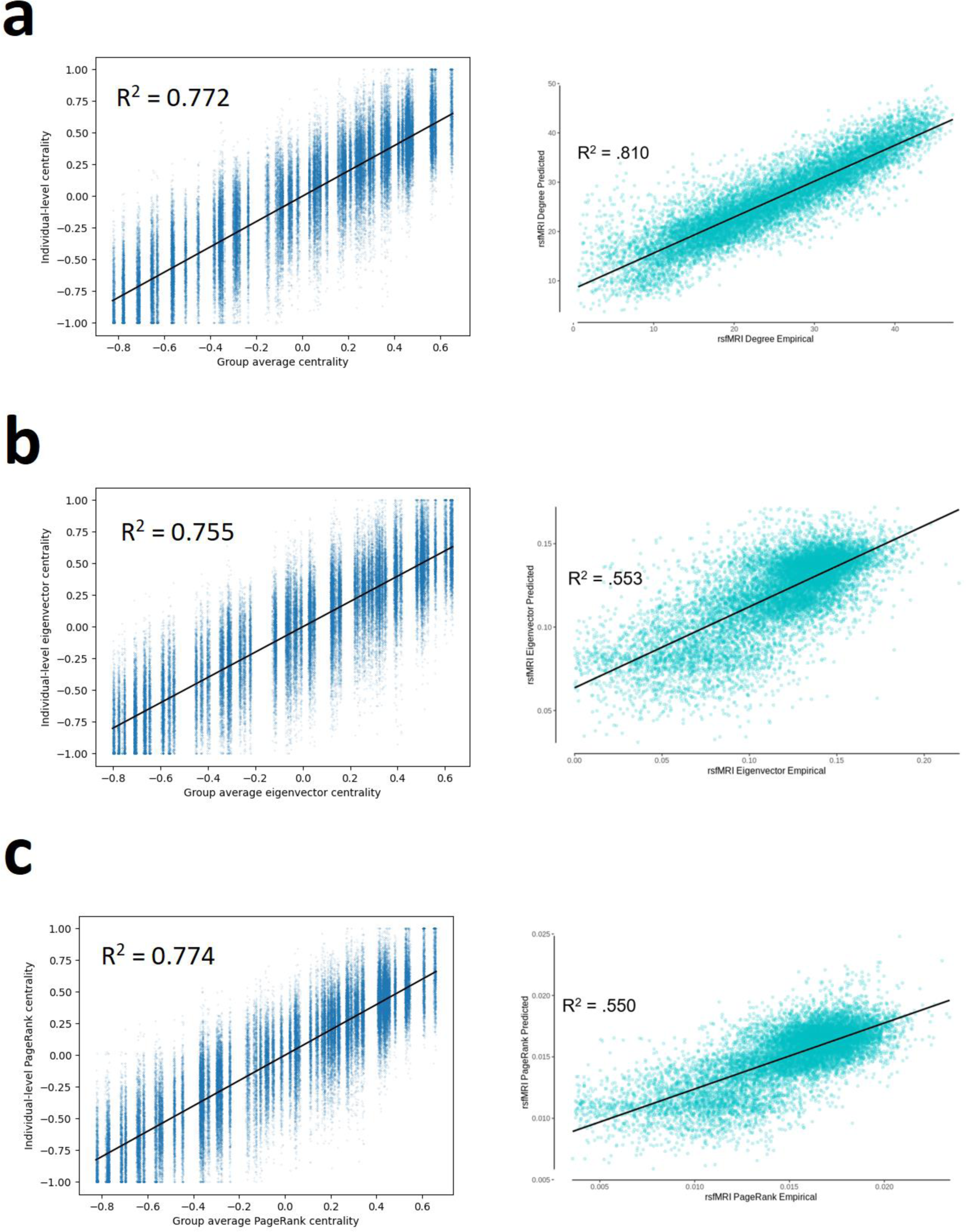
Variance in centrality measures explained by group average (left column) and pFC due to Neudorf et al. (right column) respectively for **a)** degree centrality **b)** eigenvector centrality and **c)** PageRank centrality. Right column figures adapted from (Neudorf et al., 2022), Figures S3, 5, 7 under CC-BY 4.0 license

## Discussion

Since group average FC correlates so highly with individual FC in human connectome studies, we feel that the results from individual-level structure-function prediction methods should explicitly be compared to the group average or the inherent variation in the data. Prediction methods are usually a black box from which we seek to obtain the highest possible performance, and we should compare their performance to the highest reference performance, which in this case is not a biophysical model such as considered by Sarwar et al., but rather from the group average eFC. Biophysical models enjoy the advantage of being partly explanatory, i.e. we may learn something about brain dynamics by inspecting the fitted models. Machine learning models do not typically have this property, and can thus only be judged on the usefulness of their outputs, or employ additional methods to analyze their decision-making. By extension, the comparison by Neudorf et al. to the performance of Sarwar et al. and the comparison by Benkarim et al. to other machine learning methods suffer from the same issues, since the group average is still a better predictor than all of these other methods. Deslauriers-Gauthier and colleagues also found that the group average acts as a sort of “glass ceiling” for prediction performance, considering several methods (Deslauriers-Gauthier et al., 2022). This may indicate that the current methods run into some fundamental barrier that we do not understand yet. One could argue that the performances of the prediction models provide a lower bound for the optimal performance of structure-function models, indicating that other methods such as biophysical models and network communication models could theoretically be improved to produce an individual-level eFC-pFC of at least 0.7 using the DK atlas and at least 0.77 using the Schaefer atlas. However, the group average eFC shows that we should be able to achieve at least 0.84 on the DK atlas and confirms the 0.77 lower bound for the Schaefer atlas, since any model can be “trained” to output the average training eFC when presented with any form of input data and eFC as output training data.

From these results we argue that one should consider more than just the performance of models in terms of raw correlations, since these may give an optimistic impression of prediction performance, especially when the performance of the group average is not explicitly reported in comparison. Sarwar et al. attempted this by predicting cognition, but given the results of our analyses above, it is not clear whether pFC as generated by the deep learning model is able to do so. Neudorf et al. showed that their model can explain variance in centrality measures and thus is informative of FC network structure, but the group average can do this equally well or better. Finally, Benkarim et al. infer from cortical patterns in prediction accuracy that the model captures the need for indirect connections in higher-order association areas such as the default mode and frontoparietal networks. However, we were able to show that this is simply due to FC deviating more from the group average in these regions, which might make them inherently harder to predict and does not indicate that the model has learned a meaningful aspect of SC-FC coupling.

We conclude that we cannot ascertain that the models as presented in these three studies have learned a meaningful mapping from SC to FC for the following reasons. Firstly, the group average eFC performs better in terms of correlation with individual eFC. Secondly, the individual information contained in pFC may be limited, since we could not use pFC by Sarwar et al. by itself to significantly predict cognition, and the group average performed identically on the additional analyses presented by both Neudorf et al. and Benkarim et al. Finally, the properties of noisy average FC, both in terms of inter-subject variance and prediction performance, appear almost identical to that of pFC when noise is tuned appropriately, which means that a prediction model not taking any individual information into account can mimic the results presented by each of the three studies.

### Current limitations and alternatives

It is difficult to ascertain why SC-FC predictions at an individual level in healthy subjects do not appear to be possible at this stage. It is possible that we have not yet found an appropriate model for the mapping between SC and FC, or that the machine learning models used in the three studies are not powerful enough to express such a mapping. Alternatively, it is possible that the current ways of constructing SC and FC do not capture enough relevant information to allow such a mapping. From fingerprinting studies (examining individual identifiability) we know that sufficient information *is* present to distinguish between individuals in both SC (Yeh et al., 2016) and FC (Finn et al., 2015). Then, if the data is indeed to blame, either the individual variations in SC are not informative for individual differences in functional activation patterns, or the individual differences in FC are not pronounced enough to trace them back to differences in anatomical organization.

For example, the spatial resolution of diffusion or functional MRI scans may be too coarse to capture individual differences between subjects that are informative for structure-function mapping. For resting-state functional MRI, the time resolution may also be a limiting factor, since we cannot capture individual activation pathways (which occur in the order of miliseconds), but rather only the time-delayed result of activations at 1-2 second time intervals.

Alternatively, the processing pipelines may fail to recover such differences between individuals. For instance, we may not be able to capture variation in anatomy sufficiently well using standardized atlases. There is a large variability in the sizes and anatomy of predefined regions between subjects (Caspers et al., 2006; Eichert et al., 2021; Fornito et al., 2008; Meesters et al., 2023; Ono et al., 1990). This means that normalization procedures used to register atlases onto individual scans may mislabel parts of the brain, e.g. under- or over-estimating the size of ROIs. Personalizing parcellations of the cortex by employing methods such as Bayesian networks (Kong et al., 2019, 2021) or deep learning atlases (Henschel et al., 2020; Huo et al., 2019; Wachinger et al., 2018) may alleviate these issues by ensuring that the natural variation in anatomy between individuals is accounted for. While the Desikan-Killiany atlas does some individual mapping, it may fail to capture information relevant to structure-function mapping due to being based on anatomical subdivisions, rather than connectivity.

On the structural side, the counting of streamlines to compute SC matrices may not reflect meaningful differences in connection strength, as this count will strongly depend on confounding factors such as differences in region sizes, differences in bundle diameter, and specific choices for tractography parameters (Jones, 2010; Jones et al., 2013; Smith et al., 2013; Yeh et al., 2021). Integrity measures such as Fractional Anisotropy or Mean Diffusivity may be more meaningful to quantify connection strength, although these are accompanied by some problems e.g. in and around crossing fibers (Jones, 2010; Jones et al., 2013; Vos et al., 2012). Developing metrics to measure the integrity and strength of white matter connections, e.g. building upon methods such as SIFT (Smith et al., 2015a, 2015b) or investigating alternative microstructure measures (Chamberland et al., 2019; Raffelt et al., 2012), may strongly improve the individual-level information in SC.

Regarding functional connectivity, a controversial processing step in the context of functional connectivity mapping is the definition of edges in FC. The dominant choice to calculate the Pearson correlation coefficient between the activation time series of each pair of ROIs has been extensively criticized. Criticisms regard the failure to capture time dependencies in associations between brain regions (Chang & Glover, 2010; Hutchison et al., 2013; Preti et al., 2017), the fact that Pearson correlation does not consider the causal processes that underlie activity patterns (Reid et al., 2019), or the dubious interpretability of negative correlations (Buckner et al., 2013; Murphy et al., 2009).

Several alternatives have been proposed, such as explicitly modeling functional connectivity in a dynamic way (Preti et al., 2017) or investigating the causality in activations (effective connectivity) using Granger causality (Barnett & Seth, 2014; Roebroeck et al., 2005). Although none of the alternatives has definitively proven to improve upon Pearson correlation in e.g. test-retest reliability (Noble et al., 2019), there has been little study into the structure-function relation using alternative definitions of FC. It is possible that the variation in the structure of the brain that we are able to measure with SC does not inform variation in Pearson correlation, but does inform variation in e.g. effective connectivity. Other steps in the functional preprocessing pipeline, such as the controversial Global Signal Regression (Liu et al., 2017; Mayhew et al., 2016; Murphy & Fox, 2017) or bandpass filtering (Shirer et al., 2015) could also significantly affect results. We think it is important to systematically investigate the assumptions underlying the different methods of constructing both SC and FC, and achieve some consensus on a processing pipeline that reliably captures the most informative inter-individual variation.

Finally, as in all modeling problems, an increased sample size may allow for a better fit of the proposed models and increase their generalizability.

### Future work

We believe it is important to continue this line of work, and to expand upon the methodology of the three studies in order to achieve higher prediction performance or otherwise meaningful predictions. We especially think it is important to explicitly ensure that predictive models do not simply converge to predicting some group average, such as the proposed method by Sarwar et al. did, and this practice should become the standard in structure-function prediction publications. Finally, we believe that taking a step back and re-assessing the construction of SC and FC may yield more informative versions of both, and thus improve the chances of finding a meaningful mapping from structure to function at an individual level. Specifically, for the construction of SC, more meaningful metrics of connection strength should be considered. For FC, alternatives for the Pearson correlation coefficient that take into account time dependence or causality may be more informative.

## Acknowledgements

We would like to thank prof. Andrew Zalesky and dr. Tabinda Sarwar for their valuable input and helping us to reproduce their results. The data used in this work is openly available at http://www.humanconnectome.org/. All HCP participants gave full informed consent prior to the data collection, following Washington University–University of Minnesota (WU-Minn HCP) Consortium ethical guidelines. This publication is part of the project “A personalized care path for brain tumor patients”, which is (partly) funded by The Netherlands Organisation for Health Research and Development (ZonMw), project number 10070012010006, and part of the project “Bringing Tractography into Daily Neurosurgical Practice”, which is (partly) financed by the Dutch Research Council (NWO), project number KICH1.ST03.21.004. The work of Remco van der Hofstad is supported in part by the Netherlands Organisation for Scientific Research (NWO) through the Gravitation NETWORKS grant no.\ 024.002.003. The authors declare no conflict of interest. Re-used figures were reproduced under Creative Commons (<u>CC-BY</u> and <u>CC-BY-NC-ND</u>) 4.0 license.

